# Phylogenetic analysis suggests joint control of transmission mode in a grass-endophyte symbiosis

**DOI:** 10.1101/085746

**Authors:** Alexandra Brown, Erol Akçay

## Abstract

How symbionts are transmitted between hosts is key to determining whether symbioses evolve to be harmful or beneficial. Vertical transmission favors mutualistic symbionts, and horizontal transmission more virulent ones. Transmission mode evolution itself depends on whether the host or symbiont can respond to selection on transmission mode. When hosts control the transmission mode, vertical transmission should evolve under more restrictive circumstances than when symbionts are in control. We take a phylogenetic approach to determine whether the host, symbiont, or both control transmission mode using the pooid grass-epichloid endophyte symbiosis as a model system. This study is the first to investigate control of transmission mode evolution in a phylogenetic context. We find a signal of host phylogeny but only in conjunction with symbiont identity. This pattern suggests joint control of transmission mode by the host and symbiont. It also suggests that non-genetic or non-conserved symbiont traits may determine whether host traits lead to vertical or horizontal transmission.

## 1 Introduction

Symbiotic relationships are ubiquitous and can have large impacts on the fitness of the host, symbiont, and organisms that interact with them [1, 2]. Understanding the evolution of symbiont virulence is therefore a matter of theoretical and practical interest. Transmission mode is a key factor in virulence evolution. Vertical transmission favors mutualists, and horizontal transmission parasites, assuming a positive relationship between virulence and horizontal transmission [3, 4] and in the absence of feedbacks selecting for mutualism [5, 6] or parasitism [7]. Transmission mode evolution may itself depend on whether hosts or symbionts can respond to selection on it [8]. We term this ability to respond to selection “control,” as selective pressures on the partner(s) in “control” determine the direction of transmission mode evolution. For example, in the case of parasitism, symbiont control may favor increased vertical transmission when host control does not. Despite its importance, there has been little work exploring the patterns of transmission mode evolution over the evolutionary history of extant symbioses.

In this paper, we show that a phylogenetic perspective can provide valuable insight into the control of transmission mode. In particular, we propose that if the variation in transmission mode in a given symbiosis maps onto the phylogeny of one of the partners, we can interpret it as that partner’s traits determining the transmission mode, i.e., that partner controls transmission mode. For example, if symbionts control transmission mode, related symbiont species should be more likely to employ the same transmission mode than unrelated species. If symbionts do not control transmission mode, then related symbionts should not be more likely than unrelated symbionts to employ the same transmission mode, because factors external to the symbionts determine the transmission mode. By looking at whether present-day transmission modes are correlated with host or symbiont evolutionary history, we may be able to understand which partner has controlled transmission mode evolution. In this paper, we show that this phylogenetic approach can give new insights into the question of control that can complement experimental approaches.

We study phylogenetic patterns of transmission mode in the symbiosis between cool-season grasses (subfamily Pooideae) and their fungal endophytes of the genus *Epichloë*. We chose this system because of its agricultural importance, the large amount of phylogenetic and transmission data available, and its variation in transmission mode [9, 10]. Previous work on this symbiosis has proposed host, coevolutionary, or symbiont control of transmission mode. The vertical transmission rates of asexual *Epichloë* are higher than their generally more parasitic sexual relatives, suggesting host control of transmission mode [11]. However, the fact that most *Epichloë* species are horizontally transmitted only on a related subset of their hosts suggests that host-symbiont coevolution is necessary for horizontal transmission to evolve, implying joint control of transmission mode [9]. Meijer and Leuchtmann experimentally investigated control using genetic variation in the *Brachypodium sylvaticum*-*Epichloë sylvatica* symbiosis and found that symbiont genotype correlated with transmission mode, suggesting symbiont control [12]. To our knowledge, there has not been a phylogenetic study to determine control of transmission mode across multiple symbioses.

Using recently developed methods for estimating phylogenetic effects on joint traits of interacting species [13, 14], we find phylogenetic patterns that point towards joint control of transmission mode. In particular, we find an effect of host phylogeny conditional on the symbiont’s identity. This effect, together with patterns inferred from simulated data, suggests that host traits and symbiont traits influence transmission mode, with symbiont traits evolving faster. This study is the first to investigate control of transmission mode evolution in a phylogenetic context. It points to a need for more transmission mode data to understand transmission and virulence evolution in symbioses of interest.

## 2 Methods

We determined control of transmission mode by the phylogenetic effects present in the transmission mode data. We consider five phylogenetic effects, each inducing a different correlation between host-symbiont pairs [14, 13]. The host and symbiont effects indicate host and symbiont control, respectively, and cause related hosts (respectively, symbionts) to have similar probabilities of exhibiting a given transmission mode (Figure 1a-b). The other effects indicate joint control. The coevolutionary effect (Figure 1c) causes two symbioses to be similar when both hosts and symbionts are related. It indicates that phylogenetically conserved factors in the host and symbiont interact to produce the transmission mode. The symbiont-specific host effect (Figure 1d) indicates that phylogenetically conserved host factors interact with non-conserved symbiont factors. Related hosts have similar probabilities of exhibiting a transmission mode, but these probabilities change from symbiont to symbiont regardless of symbiont relatedness. The host-specific symbiont effect (Figure 1e) arises when non-conserved host factors interact with phylogenetically conserved symbiont traits. We collected published phylogenetic and transmission data (Figure 2; supplement). We used Clann [15], Dendroscope [16], and the R [17] package APE [18], to combine the phylogenies into a supertree, removing species that appeared to have hybrid ancestry. We repeated the analysis using the two single phylogenies with the largest transmission data set. We assumed host-symbiont species pairs lacking transmission data did not form symbioses.

To simulate transmission mode data, we modified the code in [19]. We simulated each phylogenetic effect alone as well as a coevolutionary effect coupled with fast symbiont evolution. We simulated fast symbiont evolution by decreasing the correlations between related symbionts by a factor of 2 or 20. We re-analyzed simulated data with random transmission data removed to test the effect of missing data. For simulated and real data, we estimated phylogenetic effects with MCMCglmm [20] and analyzed posterior distributions with Coda [21].

**Figure 1:**
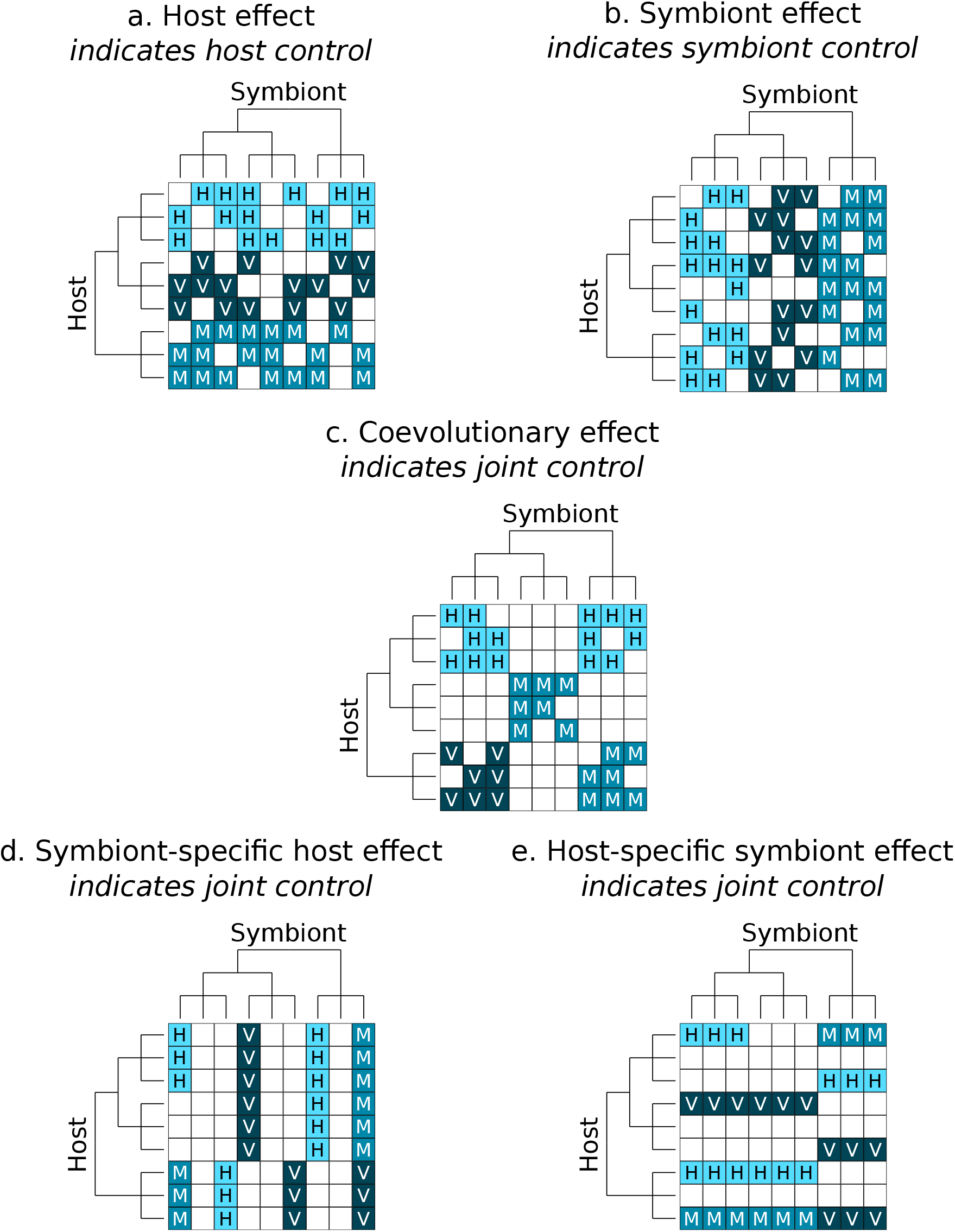
Example correlations induced by phylogenetic effects. Colored squares represent the transmission mode exhibited by host-symbiont pairs. H refers to horizontal transmission, M to mixed-mode, and V to vertical. Blank squares indicate the pair does not form a symbiosis. Host and symbiont phylogenies are shown on the left and top, respectively.

**Figure 2:**
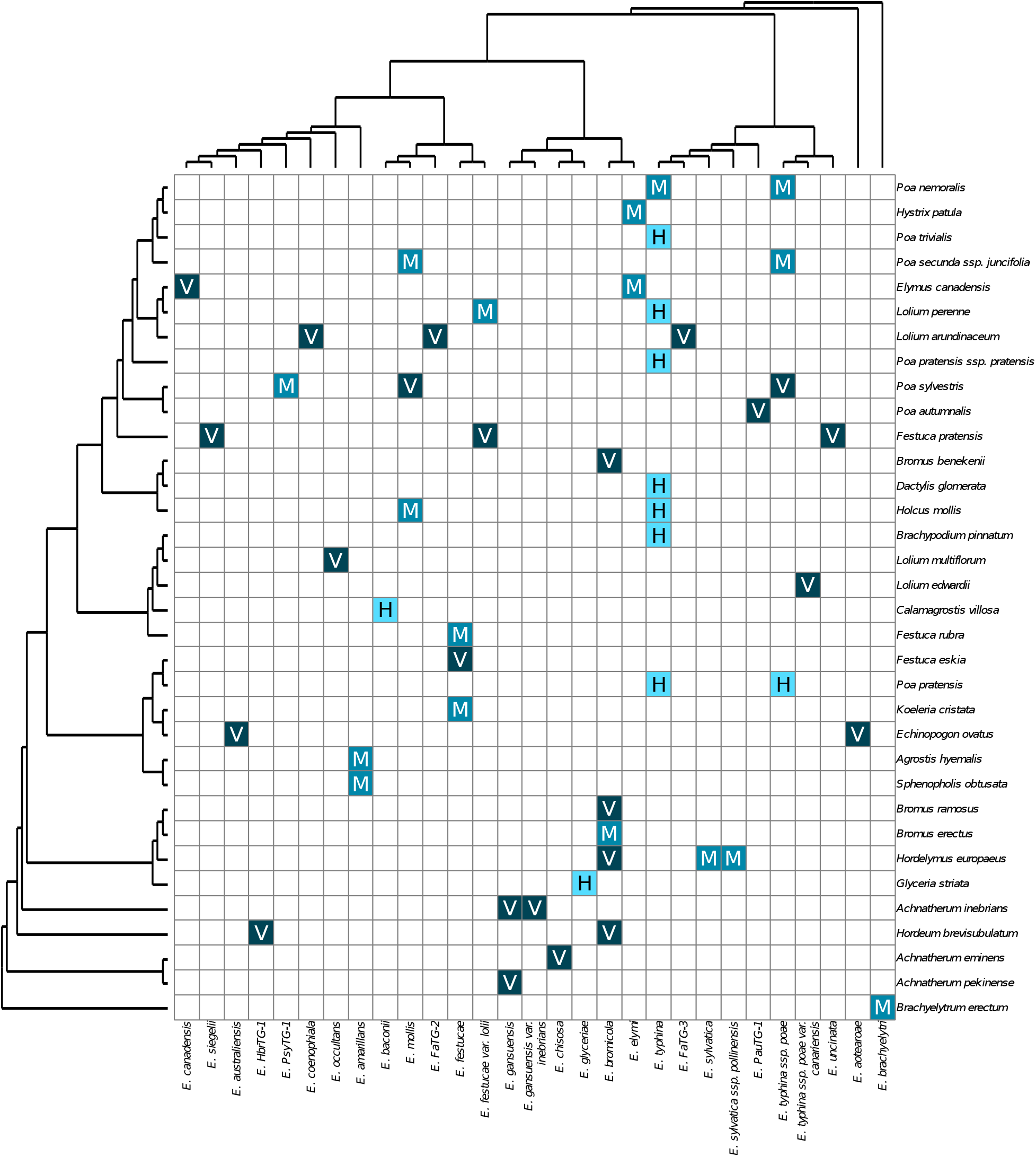
Phylogenetic and transmission mode data. Rows represent host species, columns symbiont species. Squares represent the transmission mode exhibited by a host-symbiont pair, as in Figure 1.

**Table 1:**
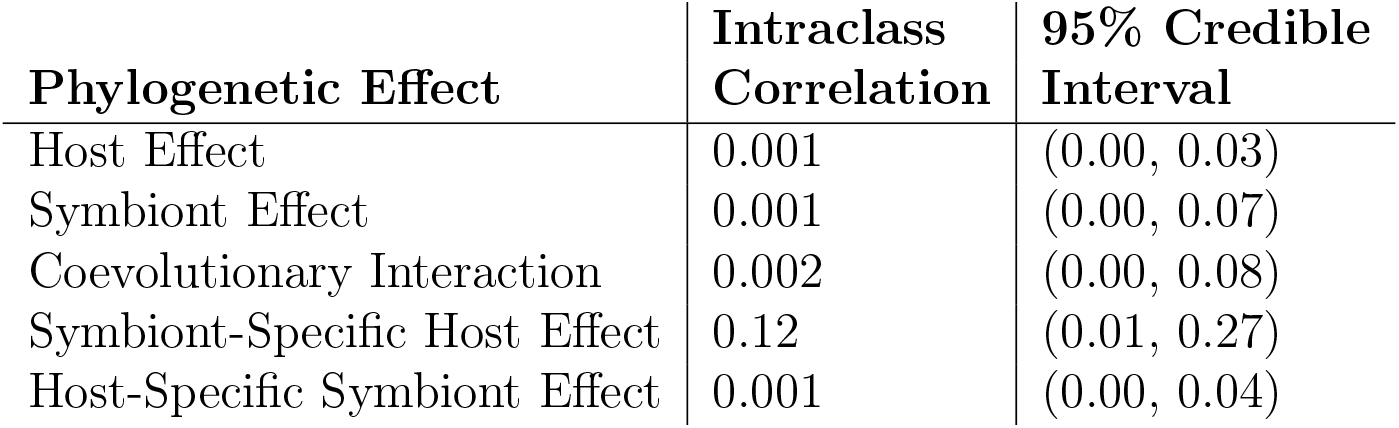
Estimated Phylogenetic Effects

| Phylogenetic Effect | Intraclass Correlation | 95% Credible Interval |
| --- | --- | --- |
| Host Effect | 0.001 | (0.00, 0.03) |
| Symbiont Effect | 0.001 | (0.00, 0.07) |
| Coevolutionary Interaction | 0.002 | (0.00, 0.08) |
| Symbiont-Specific Host Effect | 0.12 | (0.01, 0.27) |
| Host-Specific Symbiont Effect | 0.001 | (0.00, 0.04) |

## 3 Results

We detected a symbiont-specific host effect. The posterior mode of its intraclass correlation (ICC) was 0.12 (12% of the total variance in transmission mode explained), with a 95% credible interval of 1% to 27%. The host, symbiont, coevolutionary, and host-specific symbiont effects each explained no more than than 0.2% of the variance in transmission mode. The multivariate potential scale reduction factor [22] was 1.02. All effective sample sizes were > 170. When we used data from only one host and one symbiont tree, the symbiont-specific host effect explained 11% of the variance. The other effects each explained < 0.4% of the variance (Table S5).

In three out of four simulations of coevolution with fast symbiont evolution, we detected a symbiont-specific host effect. In simulations of individual phylogenetic effects at normal evolutionary rates, we generally detected the simulated effects, except for the coevolutionary effect. We detected effects that we did not simulate in seven of fifteen simulations, although in all but two simulations they appeared to be small (ICC ≤ 3%). In simulations of missing data, we never detected effects not found in the original simulated data set, although in seven of nine simulations we failed to detect effects found in the original data sets.

## 4 Discussion

We found that related hosts have similar probabilities of exhibiting a transmission mode, but their likelihood of exhibiting a particular transmission mode changes from symbiont to symbiont in a way that cannot be predicted by symbiont relatedness. This suggests that host traits interact with non-genetic or other phylogenetically non-conserved symbiont traits to determine transmission mode. Our simulation results suggest that this effect can arise from coevolutionary control of transmission mode, if rapid evolution in the symbiont masks its phylogenetic signal. Thus, it is possible that there is coevolutionary control of transmission mode in the Pooideae-*Epichloë* symbiosis. In either case, our results point to joint control of transmission mode by the host and symbiont in this symbioses.

Studies of specific grass-endophyte symbioses provide independent support for joint control. Within-species genetic variation in horizontal transmission rate has been found in symbionts in the *Pooideae*-*Epichloë* interaction [12] and in both partners in the closely related *Danthonia spicata*-*Balansia hypoxylon* symbiosis [23]. Furthermore, vertical transmission rate is phylogenetically conserved in some pooid grasses [24] and epichloid endophytes [25]. Growth rate is a possible mechanism of transmission mode control. Horizontally-transmitted endophytes outpace vertically-transmitted on certain sugars [26], while fast-growing host inflorescences can prevent symbionts from transmitting horizontally [27].

Two factors may have affected our estimates. First, some transmission data may be missing or inaccurate, given that new interactions are still being discovered [28]. Our simulations suggest that data missing completely at random rarely cause false positives (but may cause false negatives). However, non-random phylogenetic patterns in such missing interaction could change our estimates. Secondly, combining phylogenies from multiple sources may have affected our estimates of the covariance structures induced by the phylogenetic effects. This is particularly true for hybrid symbionts (e.g. *Epichloë melicicola*, which likely arose from the hybridization of the ancestors of *Epichloë aotearoae* and *Epichloë festucae*). We were only able to include hybrids when their relationship to one ancestor was missing. It is reassuring that our analysis using single phylogenies points in the same direction as the combined phylogenies.

One caveat in interpreting the phylogenetic effects is that phylogenetic effects might not map directly onto proximate control of the joint phenotype. Suppose transmission mode is proximately under host control but evolves in response to benefits provided by the symbiont. Joint control combined with high host plasticity in transmitting different symbionts may leave only symbiont phylogenetic signal detectable. Therefore, experimental work is still needed to determine proximate control. Nonetheless, quantitative phylogenetic analyses provide useful insight into how joint traits evolve.

Our results support the hypothesis that transmission mode in the Pooideae-*Epichloë* symbiosis is evolving under the control of both partners, potentially with a faster rate of evolution for the symbionts. Our analysis illustrates the potential of phylogenetic analyses in addressing questions of control in the evolutionary history of species interactions.

## Acknowledgments

We thank M. Levy for statistics advice and comments; J. Antonovics and M. Levine for comments on the manuscript; B. Koskella for comments and the scenario where plasticity masks joint control; and J. Rosario, J. Spitzer, and J. Van Cleve for computing assistance.

## Authors’ Contributions

EA and AB designed the analysis and simulations and wrote the manuscript. AB gathered data, wrote the simulations, and ran the analysis.

## Keywords

“transmission mode”,“phylogenetic effects”, *Epichloë*, “control transmission mode”

## Supplement

### 1 Methods

#### 1.1 Transmission Mode Data

We collected transmission mode data from published studies. We searched Web of Science using the following search terms: *(neotyphodium OR epichloe) AND (‘transmission mode’ OR ‘horizontal transmission’ OR ‘vertical transmission’ OR ‘mixed-mode transmission’ OR ‘mixed mode transmission’ OR ‘pleiotropic symbiosis’)*. (Asexual species in *Epichloë* were formerly in the genus *Neotyphodium*). This returned 65 papers. After discarding 18 papers whose abstracts indicated that they were unlikely to contain transmission mode data, we gathered transmission mode data from the remaining papers. We obtained transmission mode data from 32 papers [references 1–32]. 15 additional papers did not have any transmission mode data.

We recorded transmission mode as horizontal transmission, vertical transmission, mixed-mode transmission, or no transmission for each pair of host and symbiont species. A species pair was recorded as employing horizontal transmission if this was the only transmission mode reported for the pair. Similarly, vertical transmission was recorded when this was the only reported transmission mode for the species pair. A species pair was recorded as exhibiting mixed-mode transmission if it was reported to show both vertical and horizontal transmission. If no transmission data was available for a species pair, we recorded the pair as not forming a symbiosis.

Because horizontal transmission occurs via the dispersal of ascospores (although recent evidence suggests sexual reproduction is not always necessary for horizontal transmission [18, 32, 33]), a report that a symbiont was capable of reproducing sexually on a host was considered to be evidence of horizontal transmission.

#### 1.2 Phylogenetic Data

We gathered phylogenetic data from TreeBASE. We used the “All text” search option and searched for the genera of the host and symbiont species in the transmission mode data set. The search terms we used are given in Table 1. Because asexual *Epichloë* were previously members of the genus *Neotyphodium*, we also used *Neotyphodium* as a search term. Furthermore, we appended a space when searching for members of the genus *Poa* to reduce irrelevant results.

**Table 1:**
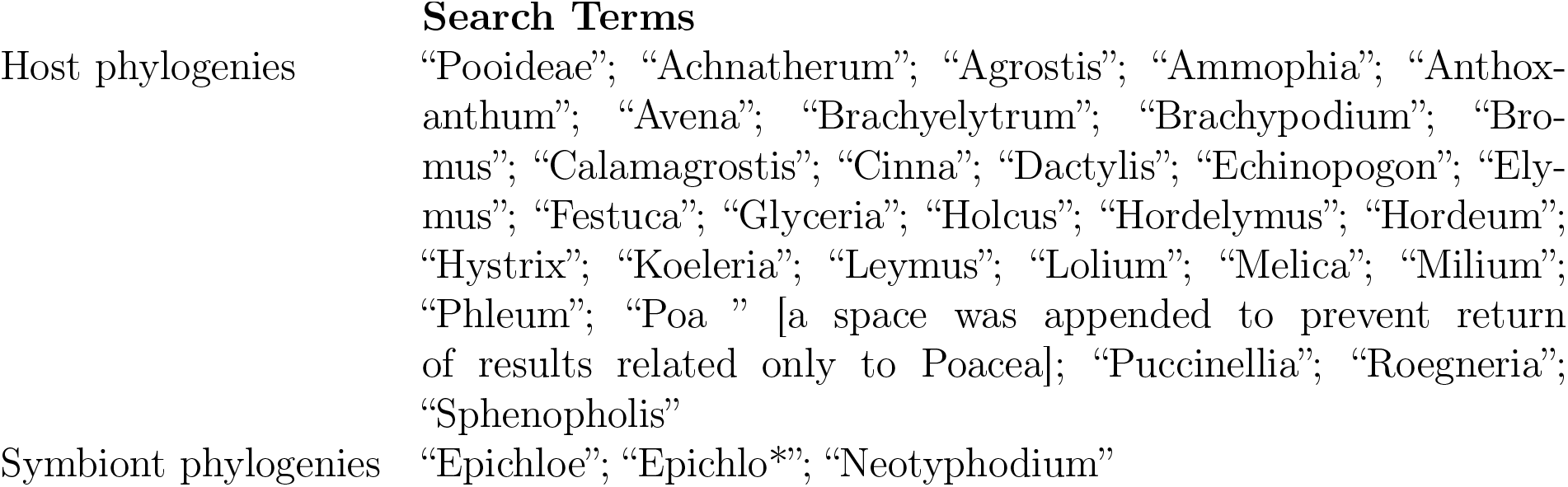
TreeBASE Search Terms

We used the R package APE [34] to remove species not present in the transmission mode data set from the trees. We deleted any trees with fewer than two species in the transmission mode data set. Because the host tree search results contained some endophyte phylogenies, we deleted any host search results that contained endophyte species. The host and symbiont phylogenies we used in the analysis are given in Tables 2 and 3, respectively.

Some phylogenies had multiple tips corresponding to the same species. We used Dendroscope’s “MUL to Network, Cluster-based” algorithm [35] to merge those species that were present twice or more in a single tree. The algorithm indicated a hybrid origin for some symbiont species. These were deleted from the trees in which they appeared to be hybrids, because we were unable to use phylogenetic networks for further analysis. To maintain as much phylogenetic information as possible, we did not delete these species from trees in which the the algorithm did not indicate a hybrid origin.

##### 1.2.1 Supertree Analysis

We used Clann [36] to find a set of equally probable host and symbiont supertreees from the trees produced merging identical tips and removing hybrids. We used the “Sub-tree Pruning and Regrafting” search algorithm, the “Most Similar Supertree” criterion, with the maximum number of steps as 3, the maximum number of swaps as 1,000,000, and 10 repetitions of the search. We used the comparisons weighting scheme and started from a neighbor-joining tree found from the average consensus distances.

Missing data were estimated using the 4 point condition distances. We combined the equally probable supertrees into a single majority consensus tree with Dendroscope’s “MUL to Network, Cluster-based” algorithm. We used these majority consensus supertrees for the main phylogenetic effects analysis.

**Table 2:** Host Phylogenetic Trees

| StudyID | TreeID | Reference |
| --- | --- | --- |
| S10359 | Tr7298, Tr7299, Tr7300, Tr7301, Tr7302, Tr7303, Tr7304 | [38] |
| S1146 | Tr1766, Tr1767, Tr1768 | [39] |
| S1304 | Tr2052 | [40] |
| S133 | Tr434 | [41] |
| S16524 | Tr78544, Tr79376 [42] |  |
| S179 | Tr4245 | [43] |
| S187 | Tr4265 | [44] |
| S2024 | Tr5125 | [37] |
| S205 | Tr4306, Tr4307, Tr4308 | [45] |
| S786 | Tr3894, Tr3895 | [46] |

##### 1.2.2 Single Tree Analysis

To test the effect of combining multiple phylogenetic trees in the analysis, we also estimated phylogenetic effects using the individual phylogenetic trees with the largest overlap with the transmission data set (Figure 1. These were T5125 for the hosts and T362 for the symbionts, both from Schardl et al. [37]. Species not present in the transmission mode data set were removed from both trees. The trees were then ultrametricized. They and the transmission data for the species in them were then used to estimate phylogenetic effects.

#### 1.3 Model of Phylogenetic Effects

In our data set, transmission mode is a categorical trait that can take on four possible values (horizontal transmission, vertical transmission, mixed-mode transmission, and no transmission). Currently, there is no method for estimating host and symbiont phylogenetic effects directly from categorical data. Thus, we modeled transmission mode for each host-symbiont pair as a 3-dimensional binary trait, following the recommendation of Hadfield and Nakagawa for estimating phylogenetic effects on categorical traits [68]. We modeled phylogenetic effects as covariances induced between the logarithms of the probabilities of each species pair expressing a given transmission mode [69, 70].

Briefly, suppose there are *n* hosts and *m* symbionts. Define the *n × m* matrix *YHT* as

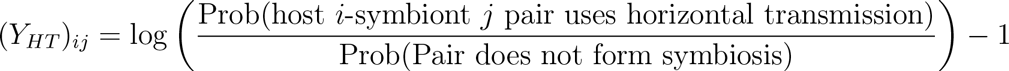

Define *Y_M M T_* and *Y_V T_* similarly for mixed-mode and vertical transmission. Then let

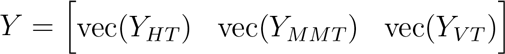

**Table 3:**
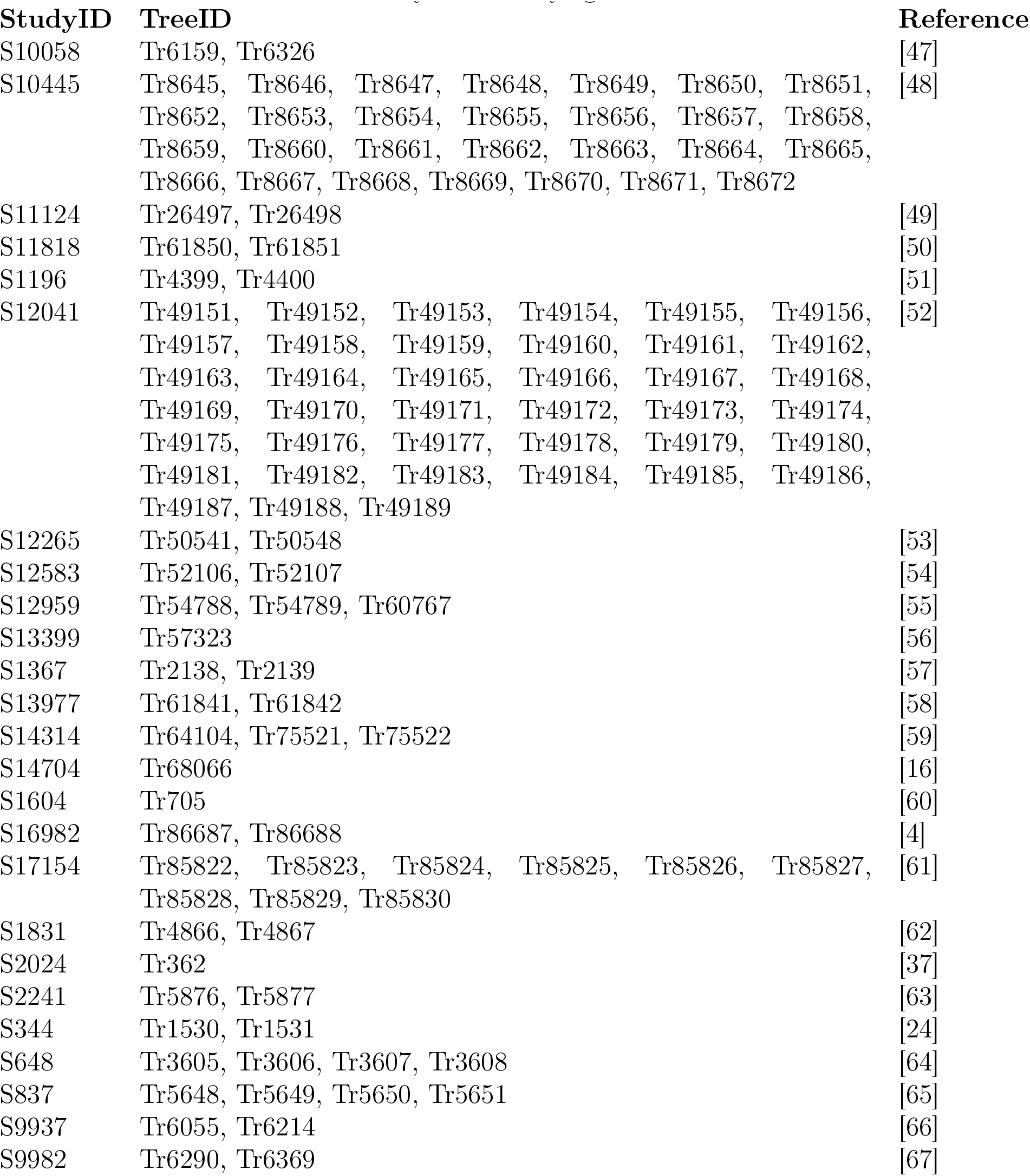
Symbiont Phylogenetic Trees

| StudyID | TreeID | Reference |
| --- | --- | --- |
| S10058 | Tr6159, Tr6326 | [47] |
| S10445 | Tr8645, Tr8646, Tr8647, Tr8648, Tr8649, Tr8650, Tr8651, Tr8652, Tr8653, Tr8654, Tr8655, Tr8656, Tr8657, Tr8658, Tr8659, Tr8660, Tr8661, Tr8662, Tr8663, Tr8664, Tr8665, Tr8666, Tr8667, Tr8668, Tr8669, Tr8670, Tr8671, Tr8672 | [48] |
| S11124 | Tr26497, Tr26498 | [49] |
| S11818 | Tr61850, Tr61851 | [50] |
| S1196 | Tr4399, Tr4400 | [51] |
| S12041 | Tr49151, Tr49152, Tr49153, Tr49154, Tr49155, Tr49156, Tr49157, Tr49158, Tr49159, Tr49160, Tr49161, Tr49162, Tr49163, Tr49164, Tr49165, Tr49166, Tr49167, Tr49168, Tr49169, Tr49170, Tr49171, Tr49172, Tr49173, Tr49174, Tr49175, Tr49176, Tr49177, Tr49178, Tr49179, Tr49180, Tr49181, Tr49182, Tr49183, Tr49184, Tr49185, Tr49186, Tr49187, Tr49188, Tr49189 | [52] |
| S12265 | Tr50541, Tr50548 | [53] |
| S12583 | Tr52106, Tr52107 | [54] |
| S12959 | Tr54788, Tr54789, Tr60767 | [55] |
| S13399 | Tr57323 | [56] |
| S1367 | Tr2138, Tr2139 | [57] |
| S13977 | Tr61841, Tr61842 | [58] |
| S14314 | Tr64104, Tr75521, Tr75522 | [59] |
| S14704 | Tr68066 | [16] |
| S1604 | Tr705 | [60] |
| S16982 | Tr86687, Tr86688 | [4] |
| S17154 | Tr85822, Tr85823, Tr85824, Tr85825, Tr85826, Tr85827, Tr85828, Tr85829, Tr85830 | [61] |
| S1831 | Tr4866, Tr4867 | [62] |
| S2024 | Tr362 | [37] |
| S2241 | Tr5876, Tr5877 | [63] |
| S344 | Tr1530, Tr1531 | [24] |
| S648 | Tr3605, Tr3606, Tr3607, Tr3608 | [64] |
| S837 | Tr5648, Tr5649, Tr5650, Tr5651 | [65] |
| S9937 | Tr6055, Tr6214 | [66] |
| S9982 | Tr6290, Tr6369 | [67] |

**Figure 1:**
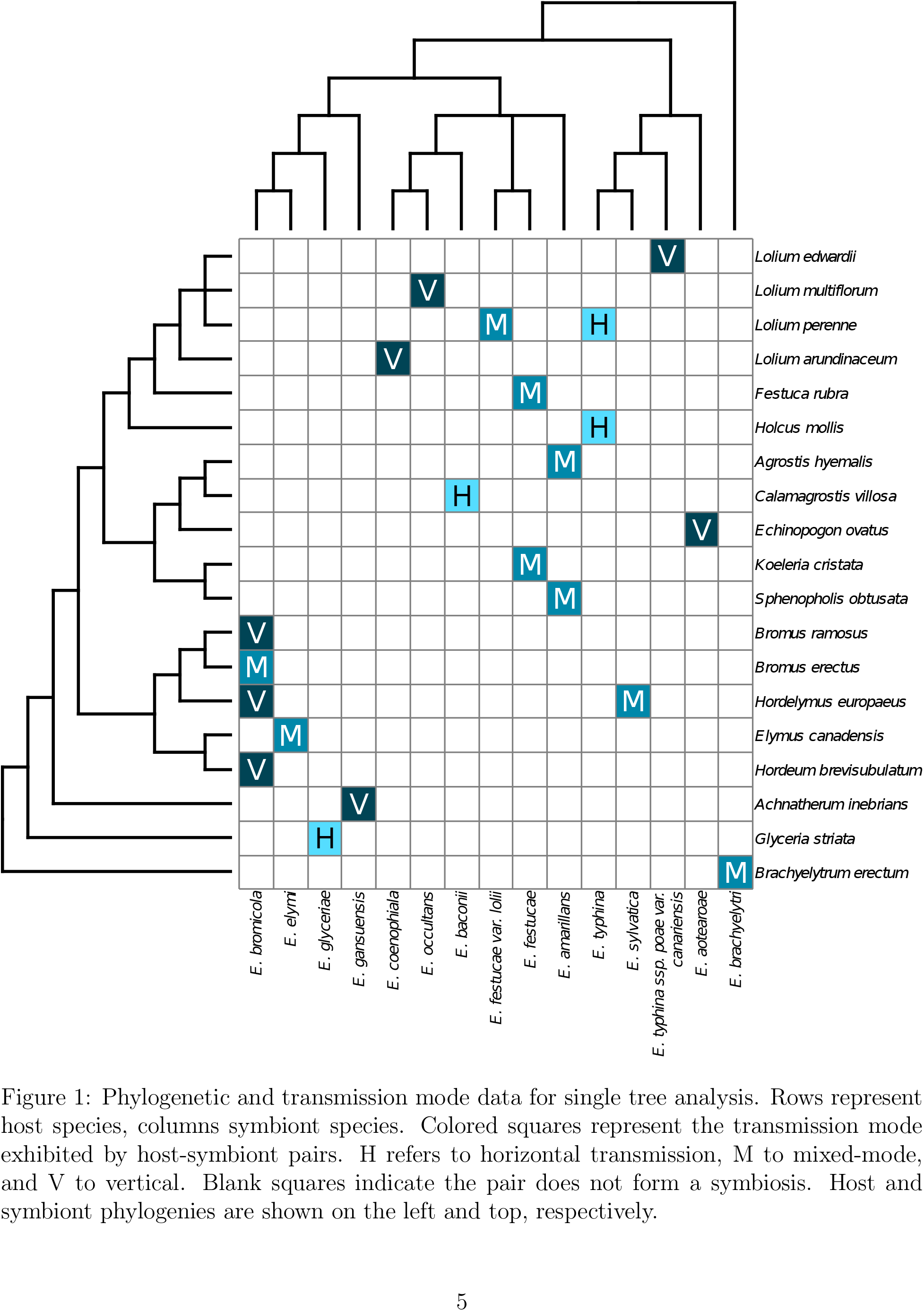
Phylogenetic and transmission mode data for single tree analysis. Rows represent host species, columns symbiont species. Colored squares represent the transmission mode exhibited by host-symbiont pairs. H refers to horizontal transmission, M to mixed-mode, and V to vertical. Blank squares indicate the pair does not form a symbiosis. Host and symbiont phylogenies are shown on the left and top, respectively.

**Table 4:**
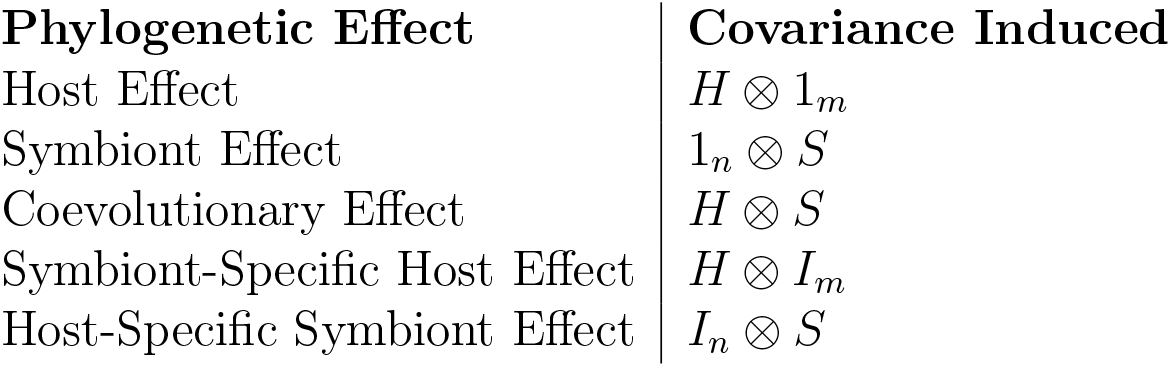
Covariances induced by each phylogenetic effect. *H* and *S* are the host and symbiont phylogenetic covariance matrices, respectively. *I_k_* is the *k × k* identity matrix. 1*_k_* is a*k × k* matrix of ones.

| Phylogenetic Effect | Covariance Induced |
| --- | --- |
| Host Effect | $H \otimes 1_m$ |
| Symbiont Effect | $1_n \otimes S$ |
| Coevolutionary Effect | $H \otimes S$ |
| Symbiont-Specific Host Effect | $H \otimes I_m$ |
| Host-Specific Symbiont Effect | $I_n \otimes S$ |

We can estimate the phylogenetic effect strengths most likely to have produced the observed transmission data if we make some assumptions about the distribution of *Y*. Specifically, we assume that *Y* has the matrix normal distribution given below, where *å*2 are the phylogenetic effect strengths to be estimated and *Vi* are the covariance structures induced by phylogenetic effects (see Table 4).

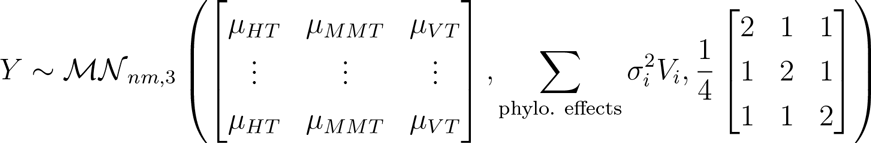

#### 1.4 Phylogenetic Effects Estimation

We estimated phylogenetic effects using the R package MCMCglmm [71]. We ran two MCMC chains for 10^6^ iterations each, with overdispersed starting values of (1) 10^-10^ for the phylogenetic effects and *-*8 for the latent variables for one chain and (2) 5 for the phylogenetic effects and *-*8 for the latent variables for the other. We used a thinning interval of 500 iterations and no burn-in. The priors for the phylogenetic effect strengths were F distributions of 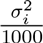 with df1 = 1, df2 = 0.002. We used the default prior for the mean, which was a multivarite normal distribution with mean 0 and variance 1 *·*10^10^ *·I*_3_, where *I*_3_ is the 3 *×*3 identity matrix.

We used slightly different analysis parameters for the single tree transmission data set because it was smaller and harder to get the MCMC chains to converge. We ran the chains for 107 iterations to allow more time for convergence. We also used a burn-in of 2000 to allow MCMCglmm to adjust the proposal distribution for the first 2000 iterations in hopes of getting a better acceptance rate. Finally, we decreased the among-column covariance (the co-variance between the log ratios of the transmission modes) from 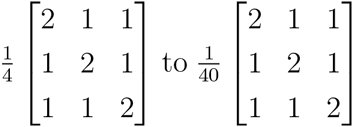 to decrease the chance of the latent variables taking on extreme values and causing numerical problems [72]. (We adjusted for this change in the among-column covariance when comparing the single tree and supertree results, discussed below.) Despite this effort to avoid numerical problems, we had to stop the analysis after 10^7^ iterations because the latent variables in one chain become too small. Fortunately, the chains appear to have converged before this point.

We rescaled our estimates of the means and calculated intraclass correlations for the phylogenetic effects, which together should have removed any difference due to differences in the among-column covariance. For all analyses, we rescaled our estimates of the means to reflect the case where the among-column covariance was 0 using the method of Diggle et al. 2002 as cited in [72]. We calculated intraclass correlations (ICCs) for the phylogenetic effects using the formula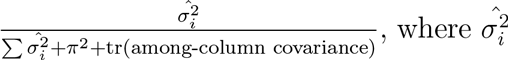 is the estimate of phylogenetic effect *i*. We did this for each saved iteration of the MCMC chains to get a posterior distribution of the ICCs.

We used the posterior mode as a point estimate of the ICC. We calculated the posterior mode using MCMCglmm’s posterior.mode function with the parameter adjust (the scaling for the bandwidth) set to 1. We obtained credible intervals for the ICCs using Coda’s HPDinterval function to get 95% highest posterior density intervals.

We checked for chain convergence using the multivariate potential scale reduction factor (MPSRF) [73] and the effective sample size, both calculated with Coda.

To analyzed our simulation results, we needed a simple way to determine whether we detected a phylogenetic effect or not. We considered an effect to have been detected if the posterior mode of its intraclass correlation was ≥0.02, or 2% of the total variance.

#### 1.5 Simulations

To determine whether the observed phylogenetic pattern can emerge from a combination of coevolutionary interactions and faster rates of evolution along the symbiont phylogeny relative to the host, we simulated transmission mode data for the case where hosts and symbionts coevolved control of transmission mode. We used the same host phylogeny used for the supertree analysis. We modified the symbiont supertree to simulate faster evolution in the symbiont. We did this by reducing the correlation between related symbionts by a factor of either 2 or 20 (but not the correlation of a symbiont with itself, which is always 1). We then simulated *Y*_sim_ as a matrix normal random variable with the variance structure given for *Y* above, with all phylogenetic effects equal to 10^−8^ except the coevolutionary effect, which we set to 4. We set the mean of *Y*_sim_ using

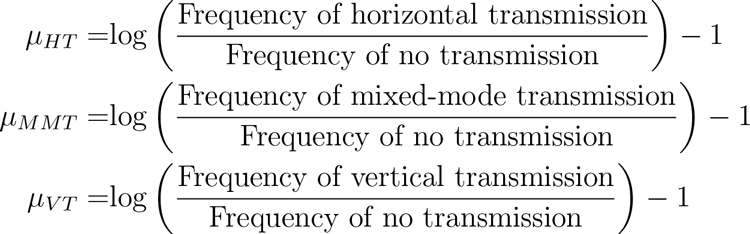

The frequencies of each transmission mode were obtained from the transmission mode data set used for the supertree analysis.

We also simulated each phylogenetic alone to test the accuracy of the phylogenetic effect estimates. For these simulations, we set one phylogenetic effect at a time to 4, and the others to 10^−8^. For each phylogenetic effect, we simulation transmission data three times. In two simulations (simulations 1 and 2 in Table 7) we simulated the case where the transmission modes were about four times as prevalent as in the dataset used for the supertree data set. Thus for *Y*_sim_ we had

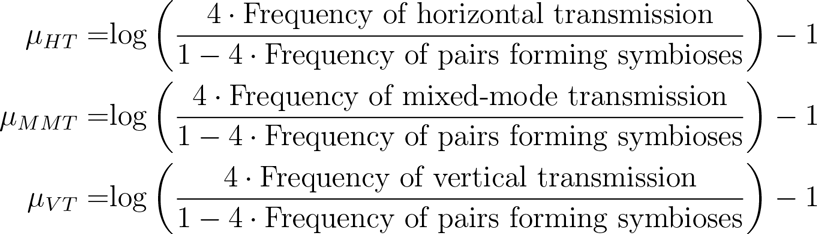
where the frequencies of transmission modes and symbioses are those in the data set used for the supertree analysis.

In the third case (simulation 3 in Table 7), we simulated the case where the transmission modes were as prevalent as in the supertree data set. In this case, *Y*_sim_ was the same as for the fast symbiont evolution simulations above.

We also simulated missing data for the simulations where the phylogenetic effects were four times as prevalent as in our supertree data set. We did this by randomly labeling 75% of host-symbiont pairs as not forming a symbiosis, whether or not they really did form a symbiosis. This meant that the simulated missing data sets had approximately the same fraction of symbioses recorded as our actual data set.

**Figure 2:**
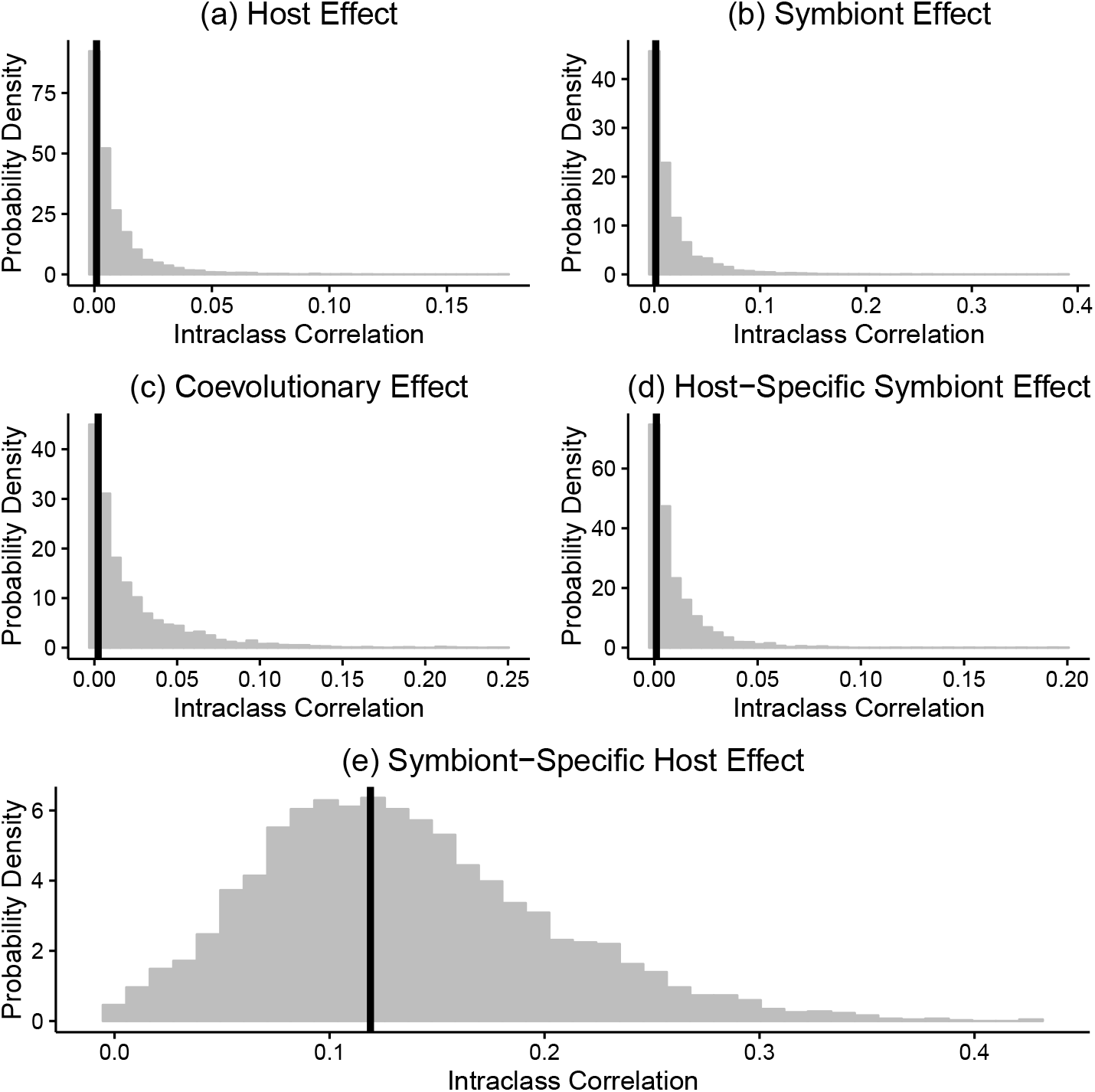
Posterior distribution of the estimates of the intraclass correlations for the supertree analysis.

### 2 Results for Analysis of Real Data

#### 2.1 Supertree Analysis Results

The posterior distribution of the intraclass correlations of the phylogenetic effects is given in Figure 2. The symbiont-specific host effect’s distribution is centered around about 12% of the total variance. The host, symbiont, coevolutionary, and host-specific symbiont effects all have most of their mass near 0% of the variance explained.

#### 2.2 Single Tree Analysis Results

Like the supertree, the symbiont-specific host effect had the only the posterior distribution with a large amount of mass on nonzero values (Figure 3). The posterior mode of the symbiont-specific host effect’s intraclass correlation was 0.11, meaning it explained 11% of the total variance, with a 95% credible interval of 0% of the total variance to 82% (Table 5). The other phylogenetic effects had posterior modes of ≤ 0.4% of the total variance.

**Table 5:**
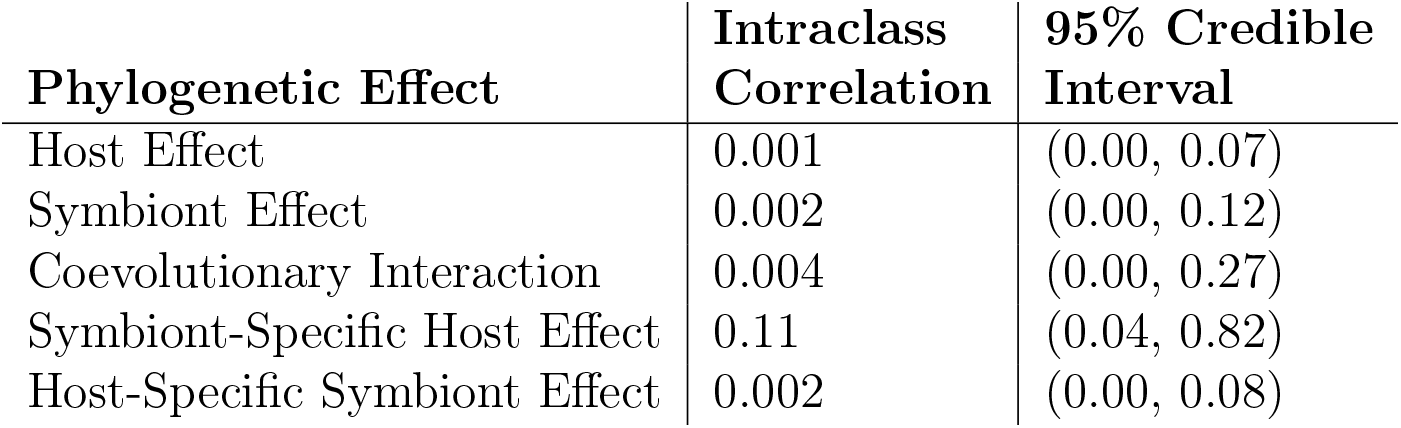
Estimated phylogenetic effects for analysis using single phylogenetic trees. Intraclass correlation given is the posterior mode.

| **Phylogenetic Effect** | **Intraclass Correlation** | **95% Credible Interval** |
| --- | --- | --- |
| Host Effect | 0.001 | (0.00, 0.07) |
| Symbiont Effect | 0.002 | (0.00, 0.12) |
| Coevolutionary Interaction | 0.004 | (0.00, 0.27) |
| Symbiont-Specific Host Effect | 0.11 | (0.04, 0.82) |
| Host-Specific Symbiont Effect | 0.002 | (0.00, 0.08) |

All the phylogenetic effects had larger credible intervals than their counterparts in the supertree data set. Further, the lower bounds of the credible intervals of the phylogenetic effects were all very close to 0% of the total variance. This may be due to the difficult of inferring phylogenetic effects on a small transmission data set.

Because of numerical problems with the MCMC chains used to estimate the phylogenetic effects for the single tree data set, we had to stop the analysis after 10 million iterations. Although the chains appeared to converge when we examined the trace plots, the multivariate potential scale reduction factor was 1.30, and the effective sample sizes of the means of log ratios of the transmission modes were in the range of 67 to 83. The symbiont-specific host effect had an effective sample size of 139. The other phylogenetic effects had effective sample sizes > 1000.

### 3 Simulation Results

#### 3.1 Fast Symbiont Evolution

Three of our four simulations of coevolution combined with fast symbiont evolution appeared to have a symbiont-specific host effect (Table 6). The posterior modes of the ICC of the symbiont specific host effect ranged from 3.8% of the total variance to 11% in these three simulations. In the fourth simulation the symbiont-specific host effect explained only 0.6% of the variance. In this simulation, the host effect had the largest ICC, explaining 3.4% of the variance. The only other phylogenetic effect detected in any simulation was a coevolutionary effect which explained 3% of the variance in one of the simulations where a symbiont-specific host effect was detected.

Based on our simulation results, it looks like it is possible for a coevolutionary effect to be mistaken for a symbiont-specific host effect when the symbiont is evolving quickly. However, we did have some difficulty in detecting a coevolutionary effect in simulations where the symbiont was evolving at the same speed as the host. It is possible that regardless of the speed of symbiont evolution, the coevolutionary effect is generally easy to mistake for a symbiont-specific host effect (which we detected in all three simulations of normal-speed coevolution below).

**Figure 3:**
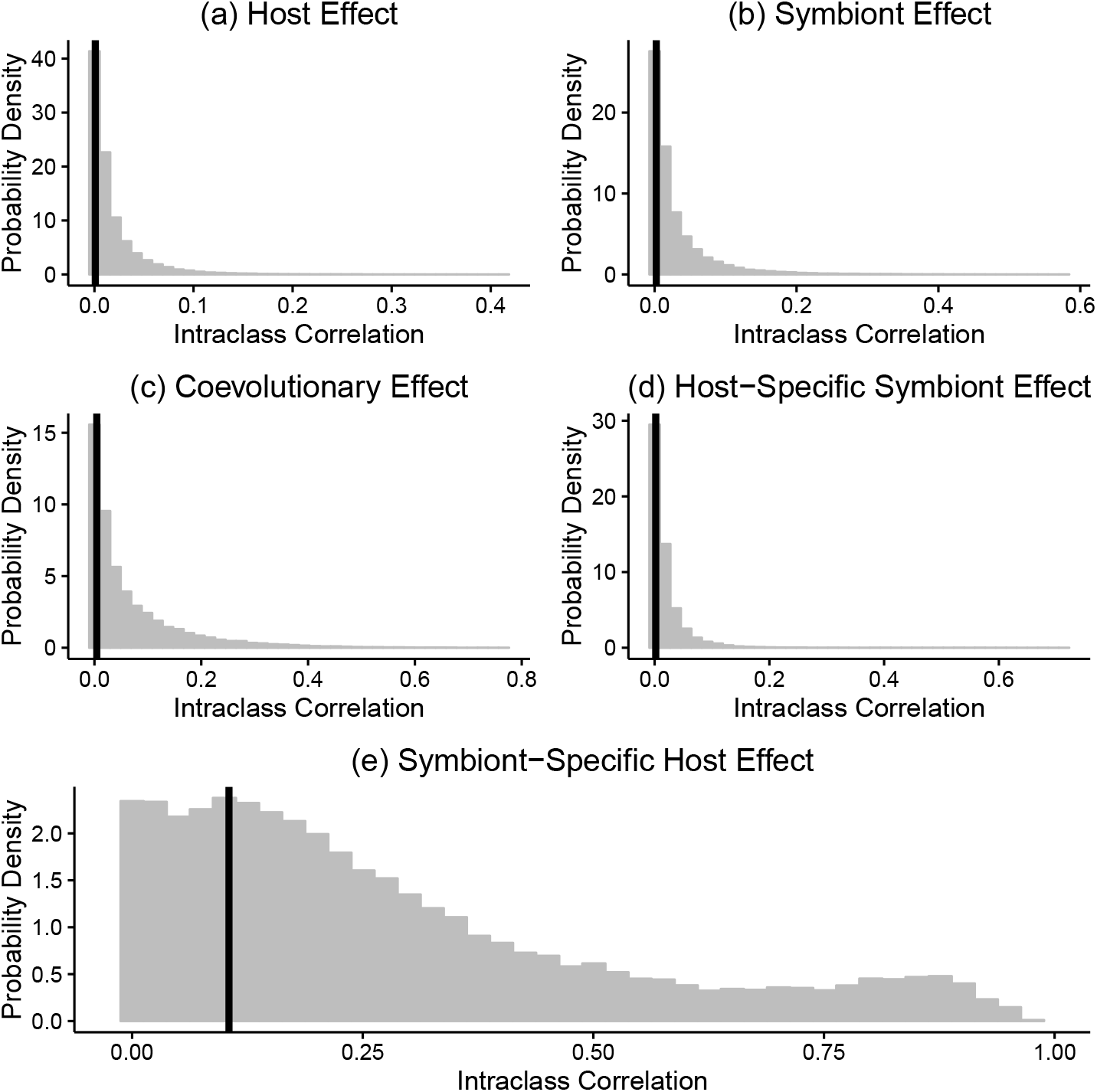
Posterior distributions of phylogenetic effects for single tree analysis.

**Table 6:** Phylogenetic effect estimates for simulated fast symbiont evolution. ICC = posterior mode of intraclass correlation.

| Symbiont<br>Evo. Rate<br>Relative<br>to Host | Simulation | Phylogenetic Effect |  |  |  |  |  |  |  |  |  |
| --- | --- | --- | --- | --- | --- | --- | --- | --- | --- | --- | --- |
|  |  | Host Effect |  | Symbiont Effect |  | Coevolutionary<br>Effect |  | Symbiont-specific<br>Host Effect |  | Host-specific<br>Symbiont Effect |  |
|  |  | ICC | 95% CI | ICC | 95% CI | ICC | 95% CI | ICC | 95% CI | ICC | 95% CI |
| 2x | 1 | **0.034** | **(0.00, 0.16)** | 0.001 | (0.00, 0.08) | 0.001 | (0.00, 0.03) | 0.006 | (0.00, 0.09) | 0.002 | (0.00, 0.05) |
| 2x | 2 | 0.001 | (0.00, 0.03) | 0.008 | (0.00, 0.13) | 0.001 | (0.00, 0.04) | **0.055** | **(0.00, 0.14)** | 0.007 | (0.00, 0.11) |
| 20x | 1 | 0.002 | (0.00, 0.06) | 0.007 | (0.00, 0.17) | **0.030** | **(0.00, 0.12)** | **0.11** | **(0.03, 0.22)** | 0.002 | (0.00, 0.05) |
| 20x | 2 | 0.001 | (0.00, 0.04) | 0.001 | (0.00, 0.05) | 0.001 | (0.00, 0.04) | **0.038** | **(0.00, 0.11)** | 0.004 | (0.00, 0.08) |

#### 3.2 Single Phylogenetic Effects

We didn’t see much difference between the data sets simulated with four times the frequency of symbioses in the real data set and those simulated with the same same frequency as in the real data. We never detected a coevolutionary effect, but otherwise we were generally successful detecting the phylogenetic effects we simulated, detecting them in at least two out of three simulations. Unfortunately, we also detected phylogenetic effects that were not present in seven of our fifteen simulated data sets (see Table 7).

When we simulated the coevolutionary effect, we detected effects other than it in three out of three simulations. In two cases we detected a symbiont-specific host effect, once in conjunction with a host effect. We detected a host-specific symbiont effect in the third simulation. Besides the coevolutionary effect simulation, there didn’t seem to be a pattern to which simulations produced false positives.

When we detected phylogenetic effects that weren’t simulated, their posterior modes were ≤ 3% in five cases. One larger effect was the symbiont-specific host effect detected in one symbiont effect simulation, which had a posterior mode of 5.8%. And in one simulation of the coevolutionary effect, a host effect was detected with posterior mode of 8.2%, and the symbiont-specific host effect had a posterior mode of 4.6%.

In all cases, the MCMC chains appeared to converge. The MPSRF was ≤ 1.04 in all analyses, and the effective sample size was ≥200.

Our simulation results suggest that we don’t have much difficulty detecting phylogenetic effects other than the coevolutionary effect, which appears to be strangely difficult to detect. However, our data set is too small for us to be certain that we will only detect phylogenetic effects that are really present in the data.

#### 3.3 Missing Data

When we simulated missing data, we never detected effects that weren’t detected in the original simulated data set (see Table 8). We failed to detect at least one effect that was detected in the original simulated data set in seven out of nine cases where at least one phylogenetic effect was detected in the original data sets. In six cases where a phylogenetic effect was detected originally, we detected no phylogenetic effects at all.

The multivariate potential scale reduction factor was ≤ 1.1 for all but one analysis, which had a MPSRF of 1.11. The effective sample size was > 100 in all analyses, and generally much larger.

Our results suggest that missing data in our real data set may cause us to fail to detect phylogenetic effects that are really present. It is possible but less likely that any missing data caused us to detect phylogenetic effects that are not really present.

**Table 7:**
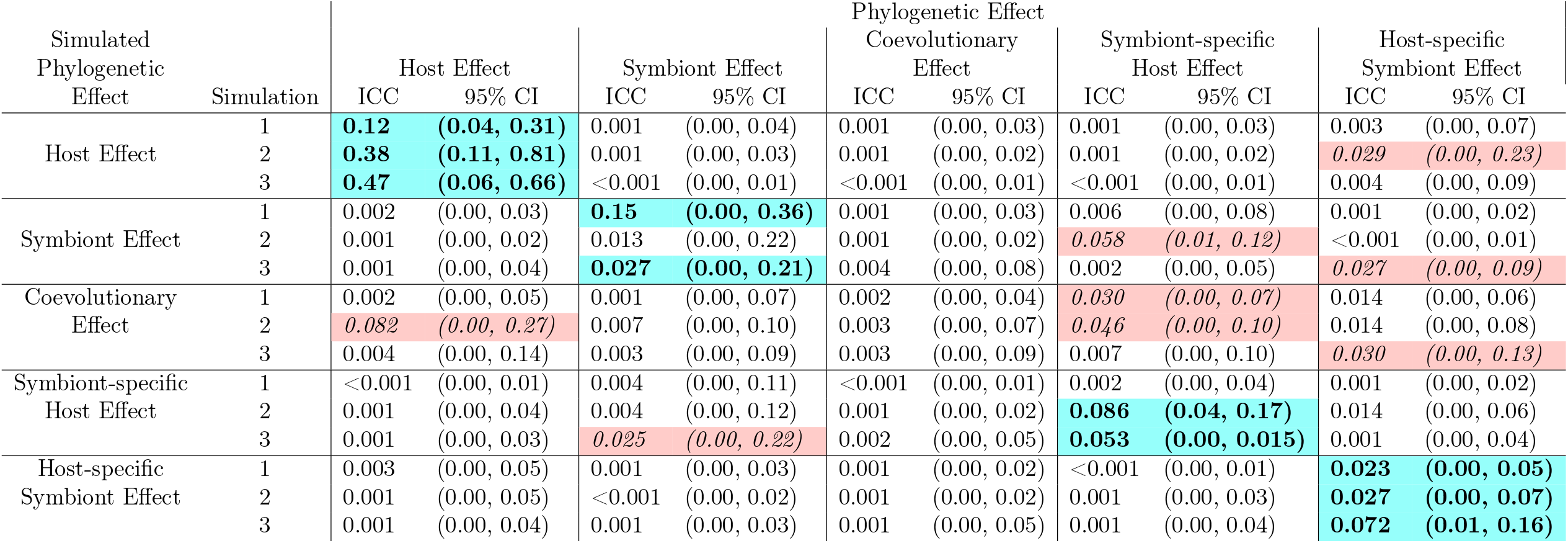
Estimated phylogenetic effects for simulated data. Colored boxes indicated phylogenetic effects that were detected (posterior mode of intraclass correlation was > 0.04). Blue boxes and bold text indicate that the effect detected was the one simulated. Red boxes and italic text indicate that the effect detected was not simulated. Simulations 1 and 2 for each phylogenetic effect had the likelihood of each transmission mode set to four times its frequency in the true supertree data set. Simulation 3 had the likelihood of each transmission mode the same as in the true data set.

**Table 8:** Estimated phylogenetic effects for simulated missing data. Blue boxes and bold text indicate that an effect detected in the original simulated data was detected (posterior mode of ICC > 0.04). Grey boxes and normal text indicate that an effect detected in the original simulated data set was not detected.

| Simulated<br>Phylogenetic<br>Effect | Simulation | Phylogenetic Effect |  |  |  |  |  |  |  |  |  |
| --- | --- | --- | --- | --- | --- | --- | --- | --- | --- | --- | --- |
|  |  | Host Effect |  | Symbiont Effect |  | Coevolutionary<br>Effect |  | Symbiont-specific<br>Host Effect |  | Host-specific<br>Symbiont Effect |  |
|  |  | ICC | 95% CI | ICC | 95% CI | ICC | 95% CI | ICC | 95% CI | ICC | 95% CI |
| Host Effect | 1 | **0.097** | **(0.00, 0.035)** | 0.002 | (0.00, 0.06) | 0.002 | (0.00, 0.07) | 0.002 | (0.00, 0.08) | 0.006 | (0.00, 0.13) |
|  | 2 | 0.006 | (0.00, 0.21) | 0.002 | (0.00, 0.08) | 0.002 | (0.00, 0.07) | 0.015 | (0.00, 0.28) | **0.029** | **(0.00, 0.27)** |
| Symbiont Effect | 1 | 0.003 | (0.00, 0.08) | **0.029** | **(0.00, 0.29)** | 0.002 | (0.00, 0.08) | 0.004 | (0.00, 0.10) | 0.011 | (0.00, 0.14) |
|  | 2 | 0.001 | (0.00, 0.03) | 0.002 | (0.00, 0.07) | 0.001 | (0.00, 0.03) | 0.004 | (0.00, 0.08) | 0.001 | (0.00, 0.03) |
| Coevolutionary<br>Effect | 1 | 0.001 | (0.00, 0.05) | 0.003 | (0.00, 0.08) | 0.001 | (0.00, 0.05) | 0.002 | (0.00, 0.06) | 0.003 | (0.00, 0.08) |
|  | 2 | 0.002 | (0.00, 0.10) | 0.005 | (0.00, 0.13) | 0.002 | (0.00, 0.07) | 0.004 | (0.00, 0.10) | 0.004 | (0.00, 0.11) |
| Symbiont-specific<br>Host Effect | 1 | 0.001 | (0.00, 0.04) | 0.003 | (0.00, 0.10) | 0.001 | (0.00, 0.04) | 0.003 | (0.00, 0.07) | 0.002 | (0.00, 0.07) |
|  | 2 | 0.001 | (0.00, 0.04) | 0.001 | (0.00, 0.07) | 0.001 | (0.00, 0.04) | 0.008 | (0.00, 0.17) | 0.008 | (0.00, 0.17) |
| Host-specific<br>Symbiont Effect | 1 | 0.002 | (0.00, 0.05) | 0.002 | (0.00, 0.07) | 0.002 | (0.00, 0.04) | 0.001 | (0.00, 0.04) | 0.007 | (0.00, 0.11) |
|  | 2 | 0.001 | (0.00, 0.06) | 0.002 | (0.00, 0.05) | 0.001 | (0.00, 0.03) | 0.001 | (0.00, 0.03) | 0.003 | (0.00, 0.07) |

